# The anti-fibrotic effects of RLN3 aggravated the pathogenesis of adolescent idiopathic scoliosis-a preliminary study

**DOI:** 10.1101/2022.03.24.485628

**Authors:** Hao Zhang, Beier Luo, Fengyuan Sun, Bing Xu, Ming Li, Guokun Wang, Zongde Yang

## Abstract

Previous research proposed that ligament laxity is a clinical feature that can be easily overlooked in patients with adolescent idiopathic scoliosis (AIS). Relaxin and relaxin-related peptides, which have anti-fibrosis roles in the vascular and kidney system and relax the pubic symphysis during pregnancy, may contribute to ligament laxity. The goal of this research was to evaluate the role of in AIS. We found significantly increased relaxin-3 plasma in patients with AIS, as well as a significant correlation between joint hypermobility and relaxin 3 plasma levels. In a classic mouse model (C57BL/6J) of scoliosis, which was established according to the literature, showed significantly higher relaxin-3 plasma levels compared to normal group. In the relaxin-3 knockout C57BL/6J mice model the prevalence of scoliosis was significantly decreased. An in vitro experiment showed that relaxin 3 has anti-fibrotic effects on spinal ligament fibroblasts in both humans and mice by inhibiting TGF-β via relaxin family peptide receptor 3 (RXFP3) and increasing expression of matrix metalloproteinase 2 and matrix metalloproteinase 9 via the TGF-β/Smad2 and MAPK-ERK1/2 pathway. Blocking RXFP3 function with R3(B1-22)R significantly decreased the prevalence of scoliosis in C57BL/6J mice. In summary, the anti-fibrotic effects of relaxin-3 and RXFP3 activation may aggravate the pathogenesis of AIS.

## Introduction

Adolescent idiopathic scoliosis (AIS) is a three-dimensional, structural deformity of the human spine that occurs in otherwise healthy children, predominantly girls. The prevalence of AIS ranges from 2%–4% worldwide^(*1, 2*)^. About 10% of those with AIS have significant deformity that requires treatment^(*3*)^. Until puberty, children with AIS show no signs of the disease^(*4*)^. Despite centuries of study and treatment, the etiopathogenesis of AIS is still not established, and there is no cure^4^. Interventions instead focus on secondary prevention to avoid negative side effects of the disease^(*5*)^. Spinal curvatures can progress to the point where surgery is needed to correct the deformity and prevent future progression. Patients often are dissatisfied with daily bracing, and they can experience back pain, alterations in body image, and pulmonary impairment as adults.

Although the etiology and pathogenesis remain unclear, ligament laxity may be a clinical manifestation of AIS and thus key to treatment and prevention of the disease. Nearly 25% of young ballet dancers have AIS^(*6*)^, which is 10 times higher than the rate of 2%–4% in the rest of the population. Other research reports that 27%^(*7*)^ and 30%^(*8*)^ of young ballet dancers have AIS, respectively. Higher rates of AIS have been found among rhythmic (12%)^(*9*)^ and other (36%)^(*10*)^ gymnasts. The research ascribed this high prevalence to joint laxity among this population, which may explain why AIS has not been observed among other athletes. Research evaluating people with AIS found that their joint mobility is significantly higher (51%) than that of a typical control group (21%)^(*11*)^, and other studies support these results(*12-14*). We believed that there may be some specific hormones lead to the ligament laxity.

Researches showed that relaxin family peptide hormones can lead to ligament laxity^(*15*)^. Relaxin is a member of a family of peptide hormones that is structurally similar to but diverged from insulin early in vertebrate evolution to form a distinct peptide family. Relaxin has a two-chain structure with a receptor binding motif and ability to bind and activate G-protein-coupled receptors(GPCRs)^(*15*)^. Relaxin alters ligament mechanics due to its collagenolytic effect, which is mediated by discharge of matrix metalloproteinases (MMPs), collagenase, and plasminogen activator^(*16*)^. However, whether relaxin contributes to the pathogenesis of AIS remains unknown. This research aimed to find a possible correlation between relaxin and AIS.

An analysis of relaxin 1, relaxin 2, and relaxin 3 plasma levels showed elevated relaxin 3 plasma levels in people with AIS and in scoliosis mice models. Furthermore, decreased prevalence of scoliosis was found in a relaxin3 knockout mouse model. Blocking function of relaxin family peptide receptor 3 (RXFP3) via R3(B1-22)R, which is an inhibitor of RXFP3^(*17-20*)^, significantly decreased the prevalence of scoliosis in a mouse model. These findings indicate that relaxin 3 has anti-fibrotic potential that may play an important role in the etiology of AIS.

## Materials and methods

### Clinical subjects

Female patients with AIS who were hospitalized at the department of spine surgery at Changhai Hospital from May 2018 to April 2020 were enrolled in this study. AIS diagnoses were made following clinical history and X-ray examination. Patients with connective tissue diseases (Marfan syndrome, Ehlers-Danlos syndrome, etc.) were excluded. The supraspinous ligament tissues were obtained from participants after posterior correction and internal fixation surgery. The plasma samples were collected participants who received brace treatment or surgical treatment. Age-matched female patients with lumbar disc herniation (without scoliosis) were enrolled as the control group. There were significant differences in body mass index, age of menarche, or Risser classification between the two groups. Table S1 summarizes the clinical characteristic of the subjects.

This study was conducted in accordance with the principles of the Declaration of Helsinki and approved by the medical ethics committee at Shanghai Changhai Hospital. Written informed consent was obtained from all participants prior to enrollment.

### Scoliosis mouse model and Rln3 knockout model

Scoliosis in mice was established via bipedal ambulation without pinealectomy, as reported previously^(*21*)^. Three-week-old C57BL/6Cnc mice were obtained from Charles River Laboratories (Beijing, China) and housed in group cages in an air-conditioned room (22 ± 2°) with controlled lighting (light from 07:00 to 19:00). During the 4 months of modeling, bipedal mice with removed forelimbs and tails were housed in special, tall cages where food and water access was progressively raised to maintain the mice’s standing postures. Scoliosis was defined as a Cobb angle >10°, according to spinal X-ray examination in the prone position. Vertebral rotation and rib hump were evaluated by helical 3D-CT. Chronic infusion of R3(B1-22)R (Phoenix Pharmaceuticals Inc, CA,USA,1μg/day)^(*19, 20*)^, and normal saline were implemented via subcutaneously implanted osmotic mini pumps (Alzet, Durect Corporation, CA, USA).

C57BL/6J mouse model with RLN3 knockout (RLN3^-/-^) was constructed by the CRISPR/Cas9 system (KOCMS181122LY1, Cyagen Biosciences Inc, Guangzhou, China; Supplemental data and Figure S1)

The medicine animal research committee of Second Military Medical University approved the animal work performed in this study. The experiment protocols were conducted according to the guidelines for the care and use of laboratory animals established by the US National Institutes of Health.

### Cell culture and identification

Human ligament fibroblasts were isolated from normal ligament tissues in the control group via type I collagenase digestion. The digested cells were maintained and passaged in Dulbecco’s Modified Eagle Medium supplemented with fetal bovine serum (10%) and penicillin/streptomycin (1%) in an incubator (37 °, 5% CO_2_). Ligament fibroblasts were identified via immunofluorescence of α-SMA. The subsequent cells experiments were performed on ligament fibroblasts at passages 3–5. TGF-β1 (10 ng/mL), RLN3 (100 ng/mL, Pepro Tech Corporation, NJ, USA)^(*18*)^, and R3(B1-22)R (0.1 μmol/L, Phoenix Pharmaceuticals Inc, CA,USA) ^(*19, 20*)^ were applied to stimulate ligament fibroblasts for 48 hours.

### ELISA assay

The plasma collected from humans and mice was subjected to ELISA assay to detect RLN concentration. Human RLN1(orb407118), RLN2(orb405219), and RLN3(orb315124) ELISA Kits were purchased from Biorbyt LLC. (St Louis, MO, USA). A Mouse Relaxin-3 ELISA Kit was purchased from Signalway Antibody LLC (MD, USA).

### Hydroxyproline measurement assay

The content of hydroxyproline in ligament fibroblasts was measured using a hydrolysate of hydroxyproline assay kit (Sigma, MO, USA) according to manufacturer instructions. Briefly, the collected cells were hydrolyzed in 100 L of concentrated hydrochloric acid (HCl, 12 M) at 120 ° for 3 hours. After centrifugation, the supernatant (10 μL) was transferred to a 96-well plate, then dried in a 60 ° oven. Each sample and standard well was successively added with chloramine T/oxidation buffer mixture and diluted DMAB reagent, then incubated for 90 minutes at 60 °. Absorbance was measured using a microplate reader (Beckman Coulter, CA, USA) at 560 nm.

### Quantitative real-time PCR

Total RNA was isolated from ligament fibroblasts using RNAiso Plus (TaKaRa Bio. Dalian, China). Equal amounts of RNA samples (200 ng) were used to generate complementary DNA using a PrimeScript RT reagent kit (TaKaRa Bio) with oligo-dT and random primers. Quantitative real-time PCR was performed on a LightCycler 480 II PCR system (Roche, Switzerland) with SYBR Green qPCR Master Mix (TaKaRa Bio). Relative gene expression levels were determined via 2-ΔΔCt analysis. Three technical replicates were established for each group. β-actin was used as a reference control.

### Immunohistochemistry assay

Protein expression and localization in ligament tissues were detected using immunohistochemistry assay, as reported previously^(*22*)^. Briefly, the deparaffinized and rehydrated sections were incubated in a warm citric acid and sodium citrate buffer (pH 6.0) for antigen retrieval. After blocking the endogenous peroxidases, the sections were incubated with the primary antibodies overnight, followed by incubation with horseradish peroxidase-conjugated secondary antibody. Color was developed using a DAB chromogen kit (Beyotime, China).

### Immunofluorescence assay

The immunofluorescence assay was performed on ligament tissues and fibroblasts, as previously described, with modifications ^(*23*)^. Briefly, the 8-μm frozen sections of ligament tissues or paraformaldehyde-fixed fibroblasts were incubated with phosphate-buffered saline containing 0.3% Triton X-100 at room temperature for 20 minutes. Then, the sections or cells were incubated with primary antibody at 4 °C overnight, followed by fluorescein-labeled secondary antibody at 37 °C for 1 hour. Nuclei were stained with 4’,6-diamidino-2-phenylindole (DAPI. Beyotime) and then visualized with a fluorescence microscope (Nikon, Japan).

### Western blot

The methods for protein purification and quantification have been described previously^(*24*)^. Protein electrophoresis separation was performed using 10% SDS-PAGE. The non-fat-milk-blocked, polyvinylidene fluoride membranes were incubated with primary antibodies overnight at 4 °C and then incubated with the secondary antibodies for 2 hours at room temperature. Visualization was achieved using ECL Plus Western Blotting Substrate (Thermo Scientific, WI, USA) on a ChemiDoc MP system (Bio-Rad, PA, USA). The relative levels of proteins were quantified based on the band grayscale value and analyzed using Image J software. β-actin was used as the internal control.

### Cell viability analysis

The ligament fibroblast growth viability was detected using a Cell Counting Kit-8 assay (Beyotime, China). The ligament fibroblasts were seeded in a 96-well plate (6 wells per group) and incubated overnight, followed by TGF-β and RLN3 treatments for 48 hours. After adding 10 μL of Cell Counting Kit-8 solution, the fibroblasts were incubated for 2 hours. The growth vitality was detected using a microplate reader (BioTek, Germany) according to the manual’s instructions.

### Statistical analysis

All statistical analyses were performed using SPSS version 22.0. The qualitative data was compared using a Fisher’s exact test. The quantitative data was compared using a two-way ANOVA or Kruskal-Wallis test when necessary. Relationships between RLN3 plasma levels and HJM points were analyzed via Pearson test. Comparison of scoliosis incidence rates was compared via chi-square test. All p values are two-sided, and values less than 0.05 were considered statistically significant.

## Results

### The elevated RLN3 level in plasma was correlated with AIS pathogenesis

To explore the function of relaxins on AIS pathogenesis, the plasma levels of RLN1, RLN2, and RLN3 were detected in patients with AIS (n=30) and healthy controls (n=30). The result from ELISA assay showed that the relaxins levels were detectable in all the human plasma samples. There was no significant difference in RLN1 level (9.71±11.08 pg/mL vs. 8.39±7.71 pg/mL; p=0.649) and RLN2 levels (76.689±50.052 pg/mL vs. 82.285±44.125 pg/mL; p=0.696) between AIS patients and healthy controls (Fig. 1A-1B). The plasma RLN3 level was significantly higher in AIS patients (84.94±64.48 ng/L) than that in healthy controls (29.42±14.26 ng/L; p<0.01; Figure 1C). Next, a nine-degree Beighton scale was applied to determine the occurrence of generalized Joint Hypermobility (JHM). The significantly higher rate was found in AIS group than that in control group (p<0.01. Supplemental Table S1). Interestingly, the plasma RLN3 level was positive correlated with the joint hypermobility in AIS patients (p<0.01. Figure 1D). Subsequently, the staining of RXFP3 (receptor of RLN3) was detected in ligament fibroblasts from both AIS and control groups by immunohistochemistry assay (Fig. 1E). Western blot assay confirmed that the expression of RXFP3 was significantly increased in the spinal ligament tissues from AIS group (Fig. 1F-1G). These results suggested that the RLN3-RXFP3 signal pathway might be close related with AIS pathogenesis. Collagen is the main matric component in ligaments and an important indicator of ligament relaxation. Therefore, by using immunofluorescence assay, the expression of collagen 1 and collagen 3 was detected in ligament tissues from patients with AIS and lumbar disc herniation. The result showed that the expression of COL1A1 and COL3A1 was significantly reduced in the spinal ligaments from AIS patients (Fig. 1H).

**Fig. 1:**
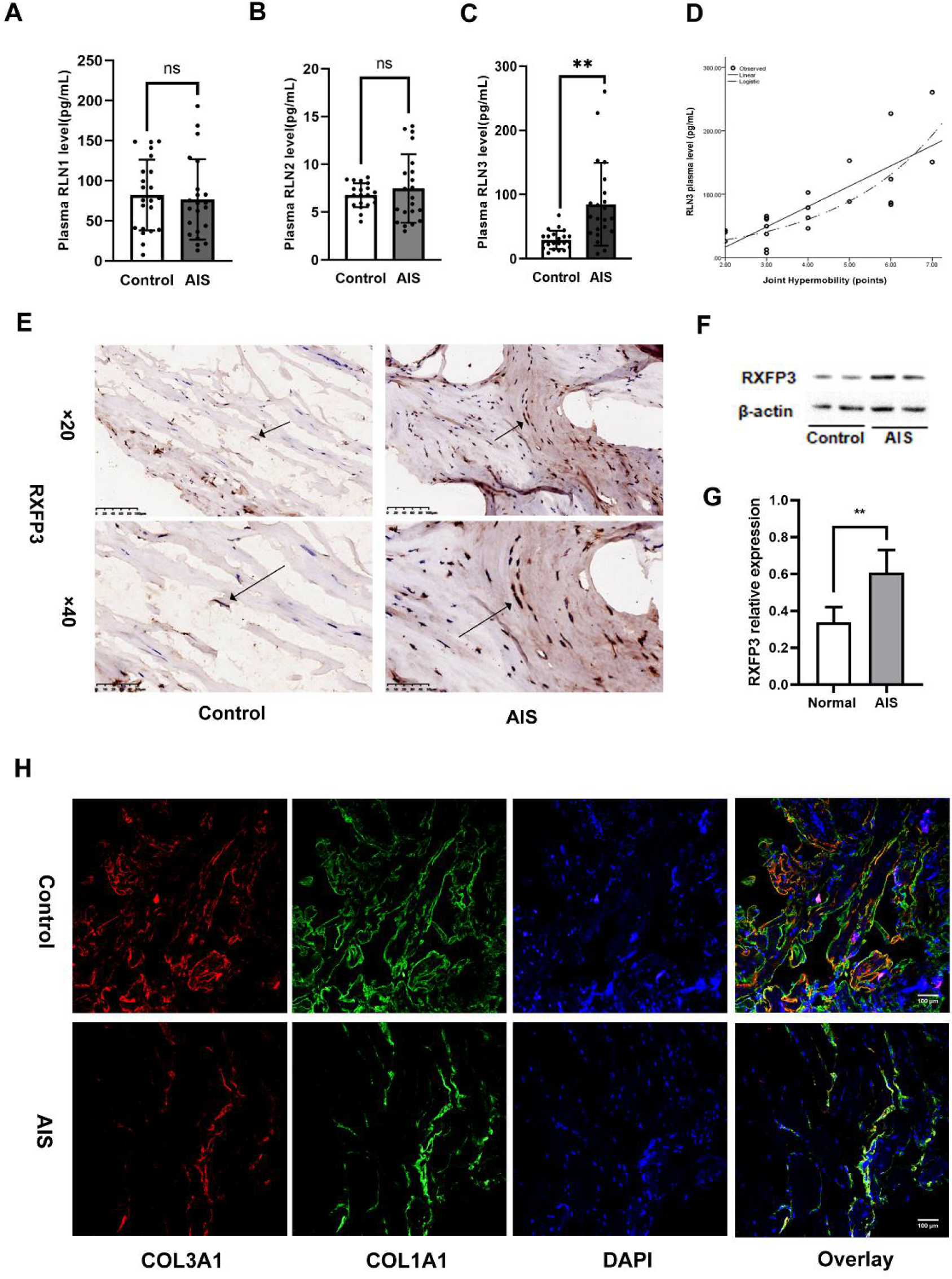
Elevated relaxin (RLN)-3 levels in the plasma RLN3 level were correlated with adolescent idiopathic scoliosis (AIS) pathogenesis. **A-C**, detection of plasma RLN1, RLN2, and RLN3 levels between the AIS (n=30) and control (n=30) groups. The plasma RLN3 level was significantly higher in the AIS group than in the control group (84.94±64.48 pg/mL vs. 29.41±14.26 pg/mL; p<0.01). No significant difference was found in RLN1 (9.71±11.08 pg/mL vs. 8.39±7.71 pg/mL; p=0.649) and RLN2 (76.689±50.052 pg/mL vs. 82.285±44.125 pg/mL; p=0.696) levels between AIS and control groups. Ns, p>0.05. **, p<0.01. **D**, The correlation analysis between joint hypermobility and the RLN3 plasma level in the AIS group (n=30), showing a significant correlation between joint hypermobility and RLN3 plasma level (p<0.01). **E**, The representative immunohistochemistry image of RXFP3 in ligament tissues from the AIS and control groups. Significantly stronger staining of RXFP3 was detected in ligaments from the AIS group. **F–G**, The expression change of RXFP3 in ligament tissues from the AIS and control groups (n=10 for each group). A western blot assay confirmed that RXFP3 expression was significantly increased in spinal ligament tissues from the AIS group. The relative protein level was evaluated based on the gray value of the corresponding band. **p<0.01. **H**, The representative immunofluorescence image of COL3A1 (Red) and COL1A1 (Green) in ligament tissues from the AIS and control groups. COL3A1 and COL1A1 were reduced in the spinal ligaments of the AIS group. DAPI (Blue) was used to stain the cell nuclei.

### RLN3-RXFP3 signal was activated during scoliosis formation

To further confirm the relationship between RLN3 and AIS pathogenesis, scoliosis animal model was established on C57BL/6J mice via bipedal ambulation ^(*21*)^. X-ray and 3D-CT examinations found that 67% (67 of 100) of C57BL/6J bipedal mice had scoliosis characterization after 4 months, including rib humps and vertebral rotation (Fig. 2A-2B). The results from ELISA assay showed that the plasma RLN3 level was significantly increased in mice with scoliosis (8.9±0.7 ng/mL), compared with those without scoliosis (7.0±0.6 ng/mL, P<0.01. Fig. 2C). Correspondingly, the increased expression of RXFP3 was also detected in the spinal ligament tissues from scoliosis mice by immunohistochemistry and western blot assays (Fig. 2D-2F). Then, the reduction of collagen 1 and collagen 3 expression was also detected in spinal ligaments from scoliosis mice by immunofluorescence assay (Fig. 2G). These results suggested that the activation of RLN3-RXFP3 signaling might be critical for scoliosis formation.

**Fig. 2:**
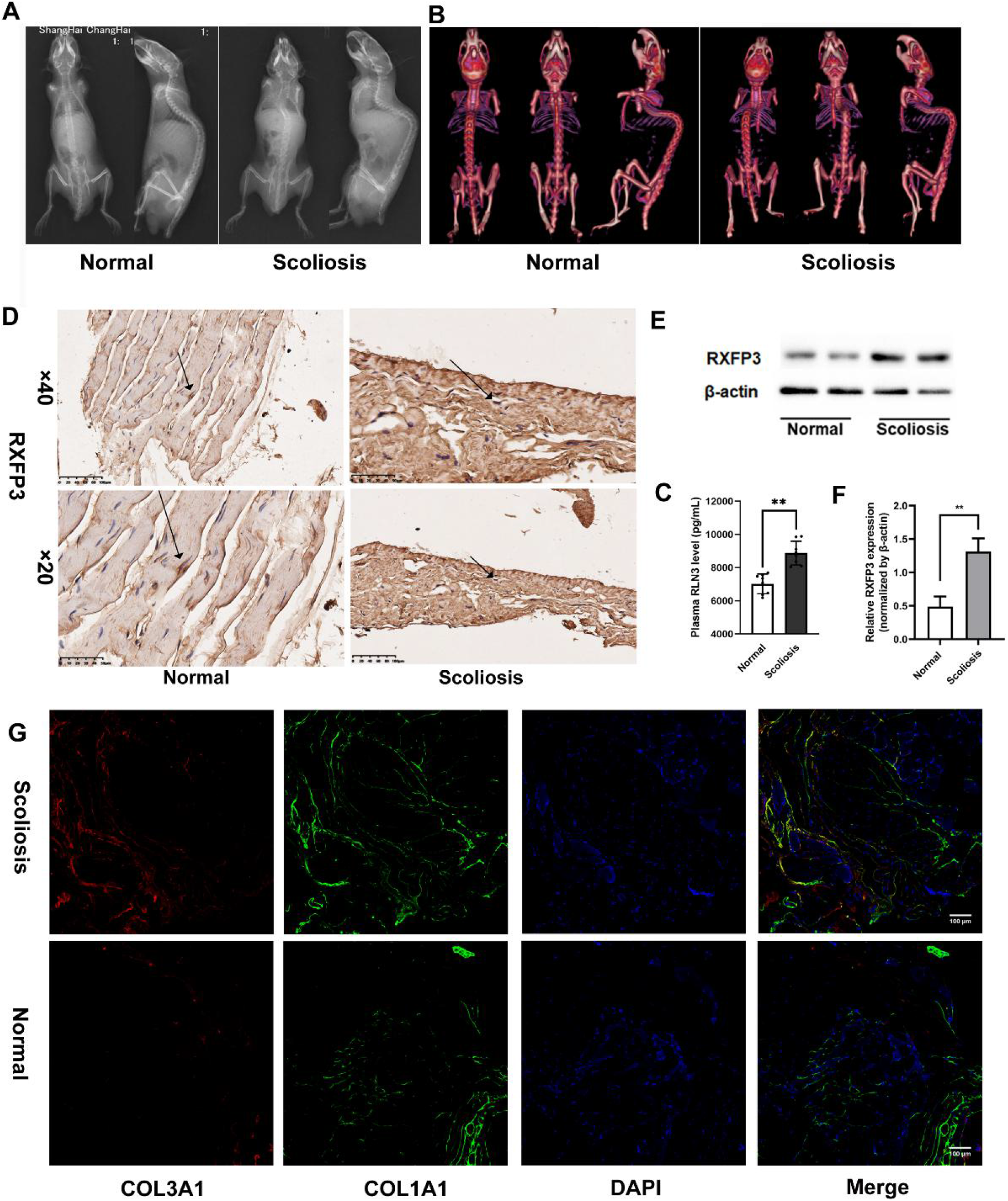
The relaxin-3 and RXFP3 signal pathway is essential for scoliosis formation in C57BL/6J bipedal mice. **A**, Scoliosis characterization in C57BL/6J mice was evaluated by X-ray. Scoliosis was found in 67% (67 of 100) of C57BL/6J bipedal mice after 4 months of feeding. **B**, 3D-CT examinations were performed on scoliosis mice. Rib humps and vertebral rotation were observed in all scoliosis mice (n=67). **C**, The relaxin-3 plasma level was evaluated in scoliosis mice (n=67) and normal mice (n=33) and found to be significantly higher in plasma from scoliosis mice (8.9±0.7 ng/mL) than in mice without scoliosis (7.0±0.6 ng/mL). **p<0.01. **D**, The representative immunohistochemistry image of RXFP3 in ligament tissues from the scoliosis and normal groups. Ligaments from scoliosis mice showed significantly stronger staining of RXFP3 than those from normal mice. **E–F**, A western blot assay confirmed significantly increased RXFP3 expression in spinal ligament tissues from scoliosis mice (n=10), compared with normal mice (n=10). The relative protein level was evaluated based on the gray value of the corresponding band. **p<0.01. **G**, The representative immunofluorescence image of COL3A1 (Red) and COL1A1 (Green) in ligament tissues from scoliosis mice and normal mice. COL3A1 and COL1A1 was reduced in the spinal ligaments of scoliosis mice. DAPI (Blue) was used to stain the cell nuclei.

### The scoliosis incidence was significantly reduced after RLN3 knockout

In order to explore the role of RLN3-RXFP3 signaling in AIS pathogenesis, a mouse model with RLN3 knockout (RLN3^-/-^) was constructed by the CRISPR/Cas9 system (Supplemental Figure S1). ELISA assay confirmed that the plasma RLN3 level was almost undetectable in RLN3^-/ -^mice (146±21pg/mL) compared with that in wild-type mice (7792±201pg/mL, P<0.0001. Figure 3C). After 4-month bipedal ambulation induction, the scoliosis incidence was significantly reduced in RLN3^-/ -^mice (19/100, 19.0%), compared with wild-type mice (64/100, 64.0%) (Figure 3A) demonstrated by X-ray and 3D-CT examinations (Figure 3B). Immunohistochemistry and western blot assays showed that the expression of RXFP3 was significantly decreased in the spinal ligament tissues from RLN3^-/-^ mice, regardless of whether scoliosis occurred (Figure 3D-3F). Immunofluorescence assay further confirmed that the decreased expression of collagen 1 and collagen 3 in spinal ligaments from scoliosis mice was significantly relieved after RLN3 knockout (Figure 3G).

**Fig. 3:**
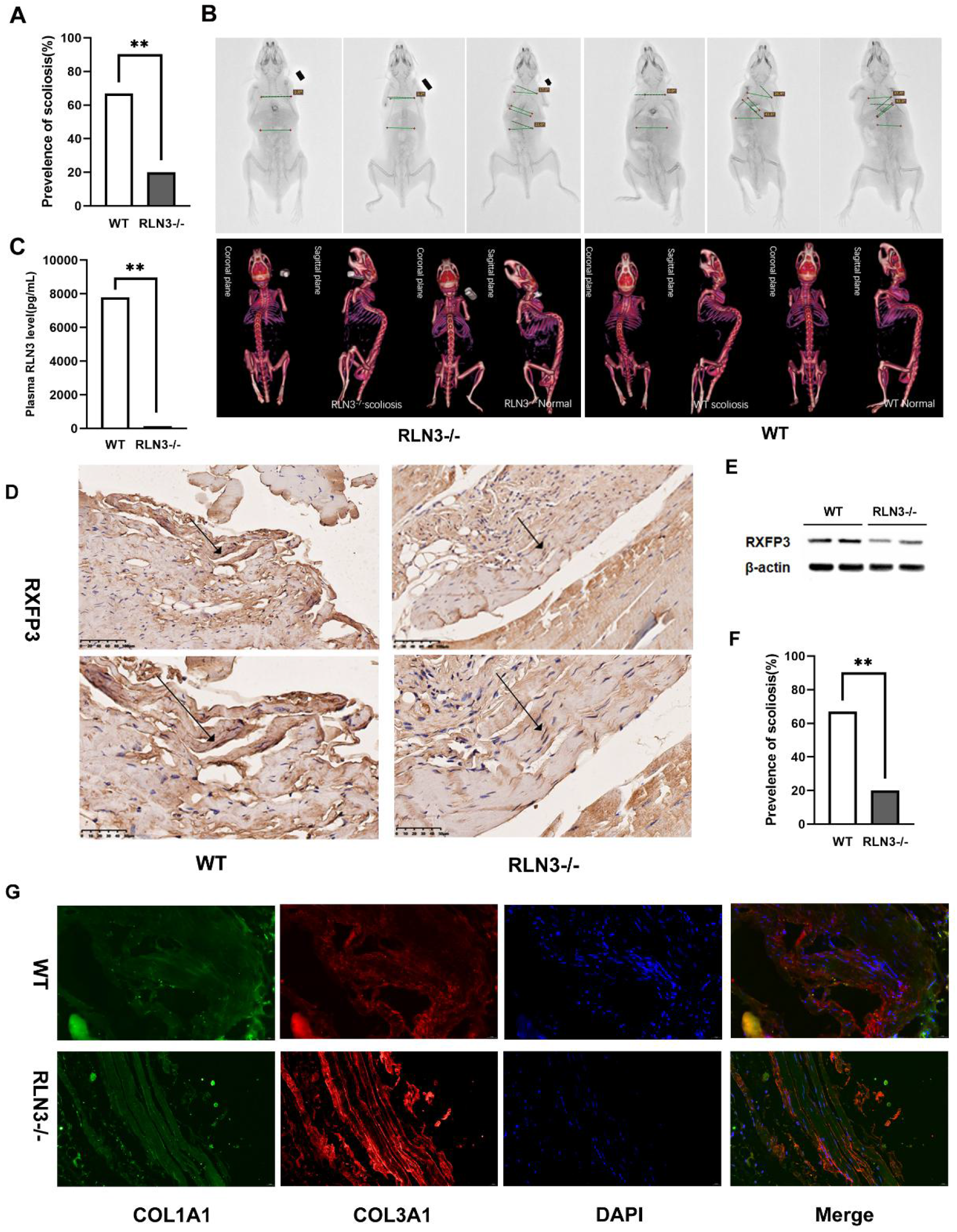
Relaxin3 (RLN3^-/-^) knockout decreased incidence of scoliosis in C57BL/6J mice. **A**, The scoliosis incidence analysis between RLN3^-/-^ and WT C57BL/6J mice. The scoliosis incidence was significantly higher in WT mice than in RLN3^-/-^mice (62% vs. 19%). **p<0.01. **B**, Scoliosis characterization in RLN3^-/-^and WT C57BL/6J mice was evaluated by X-ray and 3D-CT examinations. **C**, The detection of RLN3 levels in plasma from RLN3^-/-^mice (n=30) and WT mice (n=30). the plasma RLN3 level was almost undetectable in RLN3^-/-^mice (146±21pg/mL) compared with that in wild-type mice (7792±201pg/mL), **p<0.001. **D**, The representative immunohistochemistry image of RXFP3 in ligament tissues from the RLN3^-/-^and WT groups. Low expression of RXFP3 was detected in the spinal ligaments from all RLN3^-/-^mice, including the scoliosis and normal mice. **E–F**, Western blots confirmed that the expression of RXFP3 was significantly decreased in the spinal ligament tissues from RLN3^-/-^mice (n=10), regardless of whether scoliosis occurred. The relative protein level was evaluated based on the gray value of the corresponding band. **p<0.01. **G**, The representative immunofluorescence image of COL1A1 (Green) and COL3A1 (Red) in ligament tissues from RLN3^-/-^mice and WT mice. Relieved COL1A1 and COL3A1 was observed in the spinal ligament tissues of RLN3^-/-^scoliosis mice. DAPI (Blue) was used to stain the cell nuclei.

### RLN3-RXFP3 inhibited ligament fibroblast activation but promoted MMP2 and MMP9 expression

Activation and enrichment of fibroblasts induces sustained secretion of matrix proteins, leading to extracellular matrix remodeling. Therefore, we investigated the degree of fibroblast activation in ligaments from patients with AIS and from healthy controls. An immunohistochemistry assay confirmed decreased signaling of α-SMA in ligaments from patients with AIS (Fig. 4A). A western blot assay found significantly lower expression of α-SMA in spinal ligaments from the AIS group than those from the control group (Fig. 4B and 4C). These results suggest that activation of ligament fibroblasts was significantly inhibited during AIS pathogenesis.

**Fig. 4:**
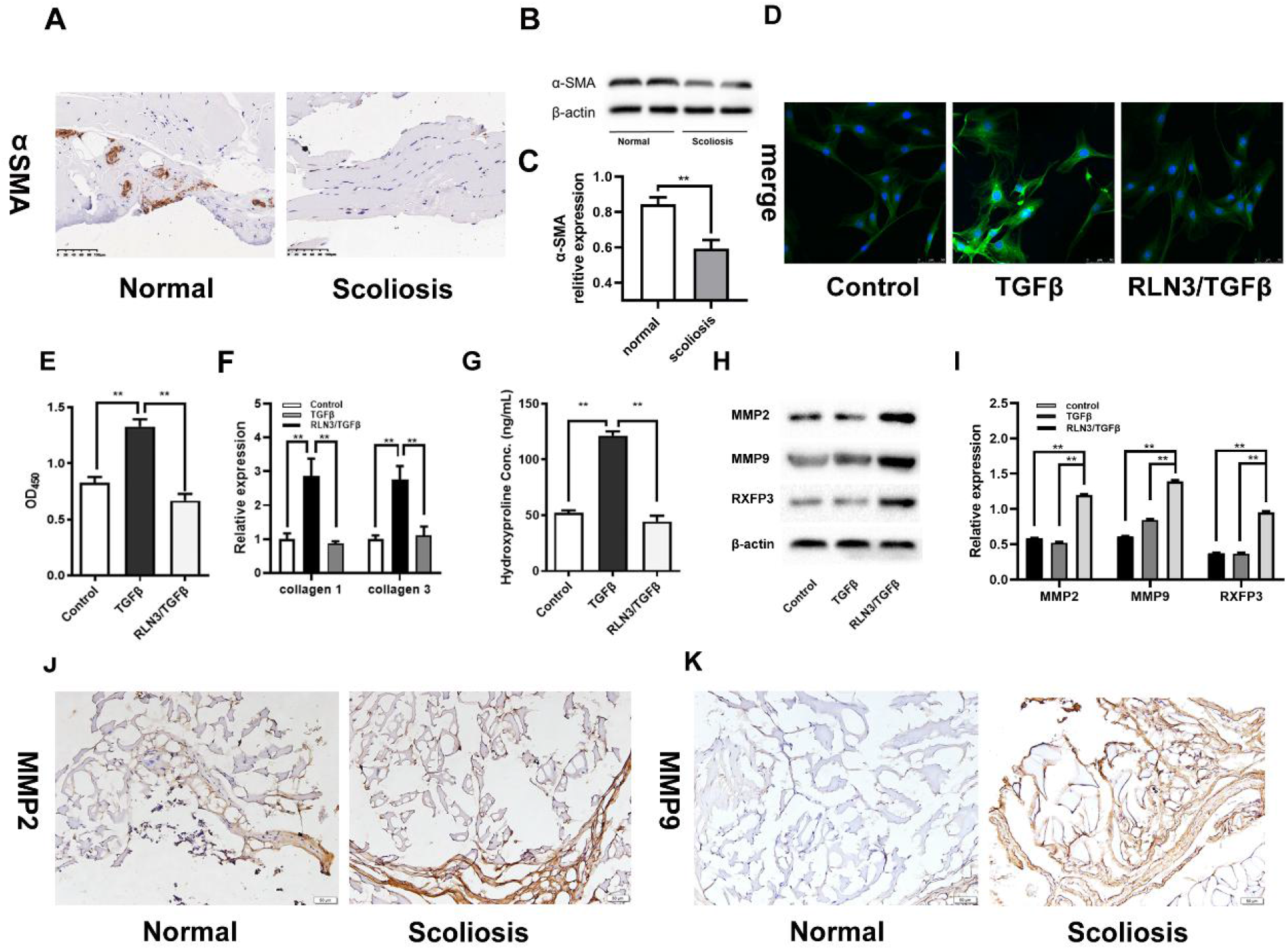
Relaxin-3 and RXFP3 activation inhibited spinal ligament fibroblast activation but promoted MMP2 and MMP9 expression. **A**, The representative immunohistochemistry image of α-SMA in spinal ligament tissues from the AIS and control groups. High expression of α-SMA was detected in the control group. **B–C**, Western blot assays found significantly lower expression of α-SMA in spinal ligament tissues from the AIS group (n=5) than those from the control group (n=5). The relative protein level was evaluated based on the gray value of the corresponding band. **p<0.01. **D**, The representative immunofluorescence image of α-SMA in human spinal ligament fibroblasts treated by TGF-β1 (10 ng/mL) and RLN3 (100 ng/mL). The expression of α-SMA was activated by TGF-β1 in human spinal ligament fibroblasts but inhibited by RLN3 treatment. **E**, A CCK-8 assay confirmed that the activating role of TGF-β1 (10 ng/mL) on growth vitality of human spinal ligament fibroblasts was significantly inhibited by RLN3 (100 ng/mL) treatment. **p<0.01. **F**, A QRT-PCR assay confirmed notably elevated expression of COL1A1 and COL3A1 in TGFβ1-treated spinal ligament fibroblasts, which was neutralized by RLN3 treatment. **p<0.01. **G**, The content of hydroxyproline in ligament fibroblasts was measured using a hydrolysate of hydroxyproline assay kit. Hydroxyproline was significantly increased in TGFβ1-treated ligament fibroblasts but released by RLN3 treatment. **p<0.01. **H–I**, Western blot assays confirmed significantly increased expression of MMP2, MMP9, and RXFP3 in spinal ligament fibroblasts after RLN3 treatment. The relative protein level was evaluated based on the gray value of the corresponding band. **p<0.01. **J–K**, The representative immunohistochemistry images of MMP2 and MMP9 in ligament tissues from the scoliosis and normal groups. Elevated expression of MMP2 and MMP9 was detected in ligament tissues from scoliosis mice.

Then, we tested the role of RLN3 on proliferation and activation of human ligament fibroblasts. The activation and growth vitality of fibroblasts were significantly inhabited by RLN3 treatment, which was confirmed by immunofluorescence of α-SMA and Cell Counting Kit-8 assays (Fig. 4D and 4E). Meanwhile, the expression of collagen 1 and collagen 3 decreased in RLN3-treated fibroblasts (Fig. 4F). An ELISA assay also showed reduced hydroxyproline content in the culture supernatant from the RLN3-treated group (Fig. 4G). Western blot further confirmed that the expression of MMP2 and MMP9 was significantly increased in fibroblasts after RLN3-RXFP1 activation (Fig. 4H and 4I). Using immunohistochemistry assay, we detected elevated expression of MMP2 and MMP9 in ligament tissues from scoliosis mice (Fig. 4J and 4K).

### RLN3-RXFP3 activated ERK1/2 signaling but inhibited TGF-β/SMAD signaling

To explore the downstream molecular mechanism of RLN3-RXFP3 signaling during AIS pathogenesis, we further detected the expression of fibroblast activation-related signaling pathways in ligament fibroblasts in response to RLN3 stimulation. A western blot assay proved that RLN3-RXFP3 could activate ERK1/2 signaling in ligament fibroblasts, demonstrated by the increased phosphorylation level of ERK1/2 and the high expression of neuronal nitric oxide synthase in response to RLN3 stimulation. Meanwhile, TGF-β/SMAD signaling was significantly inhibited by RLN3-RXFP3, demonstrated by the decreased phosphorylation level of SMAD2 (Fig. 5A and 5B). Predictably, these changes in ligament fibroblasts induced by RLN3 could be significantly neutralized by R3(B1-22)R, an inhibitor of RXFP3 (Fig. 5C and 5D) ^(*25*)^.

**Fig. 5:**
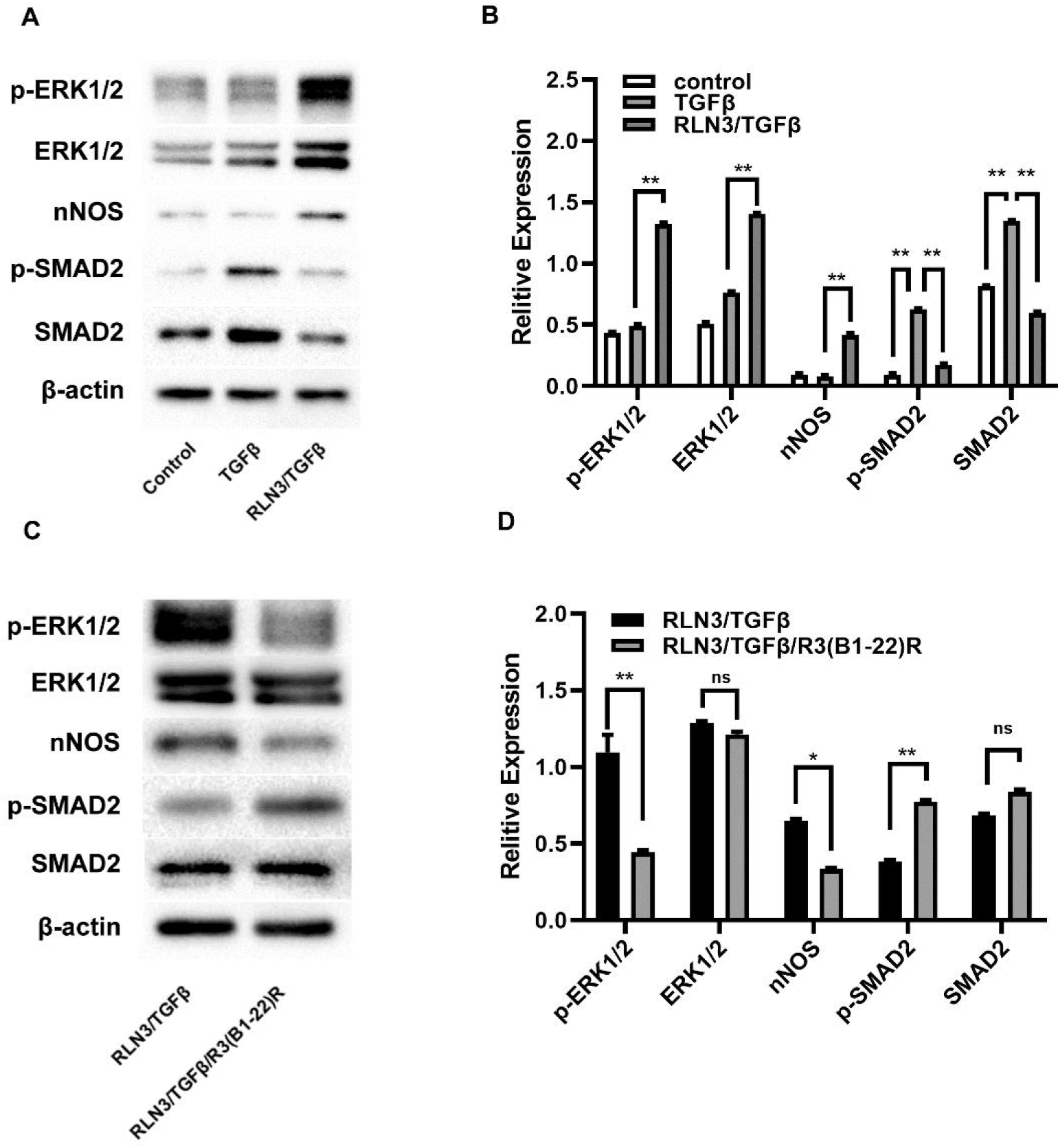
Relaxin-3 (RLN3) and RXFP3 activated ERK1/2 signaling but inhibited TGF-β/SMAD signaling. **A–B**, The role of RLN3 treatment on ERK1/2 and TGF-β/SMAD signaling in human spinal ligament fibroblasts. Western blot assays confirmed that RLN3 stimulation activated expression of phosphorylated ERK1/2 and neuronal nitric oxide synthase in ligament fibroblasts and decreased expression of phosphorylated SMAD2. Three biological replicates were set for each group. The relative protein level was evaluated based on gray value of the corresponding band. **p<0.01. **C–D**, The role of R3(B1-22)R (an inhibitor of RXFP3) on ERK1/2 and TGF-β/SMAD signaling in human spinal ligament fibroblasts. Western blot assays confirmed that R3(B1-22)R significantly neutralized RLN3-induced phosphorylation of ERK1/2 and increased expression of neuronal nitric oxide synthase. Three biological replicates were set for each group. The relative protein level was evaluated based on the gray value of the corresponding band. *p<0.05, and **p<0.01.

### Inhibition of RLN3-RXFP3 by R3(B1-22)R prevented scoliosis in C57BL/6J mice

To further explore the potential of RLN3-RXFP3 as a therapeutic target for AIS, the R3(B1-22)R inhibitor^(*25*)^ was administered to C57BL/6J mice with scoliosis. X-ray and 3D-CT examinations (Fig. 6A and 6B) confirmed that the incidence of scoliosis in the R3(B1-22)R-treated group was significantly lower than that of the NS group (22.0% vs. 62.0%), as shown in Fig. 6C. Compared with spinal ligaments from normal mice (Fig. 2G), the expression of collagen 1 and collagen 3 significantly increased in most mice treated with R3(B1-22)R and significantly decreased in most mice treated with normal saline (Fig. 6D). Meanwhile, the expression of MMP2 and MMP9 significantly decreased in the spinal ligaments, respectively, of mice treated with R3(B1-22)R (Fig. 6E and 6F). However, there was no obvious RXFP3 expression change in spinal ligaments between the NS and R3(B1-22)R groups (Fig. 6G). Moreover, a western blot assay confirmed that treatment with R3(B1-22)R could significantly inhibit ERK1/2 phosphorylation and neuronal nitric oxide synthase expression and promote SMAD2 phosphorylation (Fig. 6H and 6I).

**Fig. 6:**
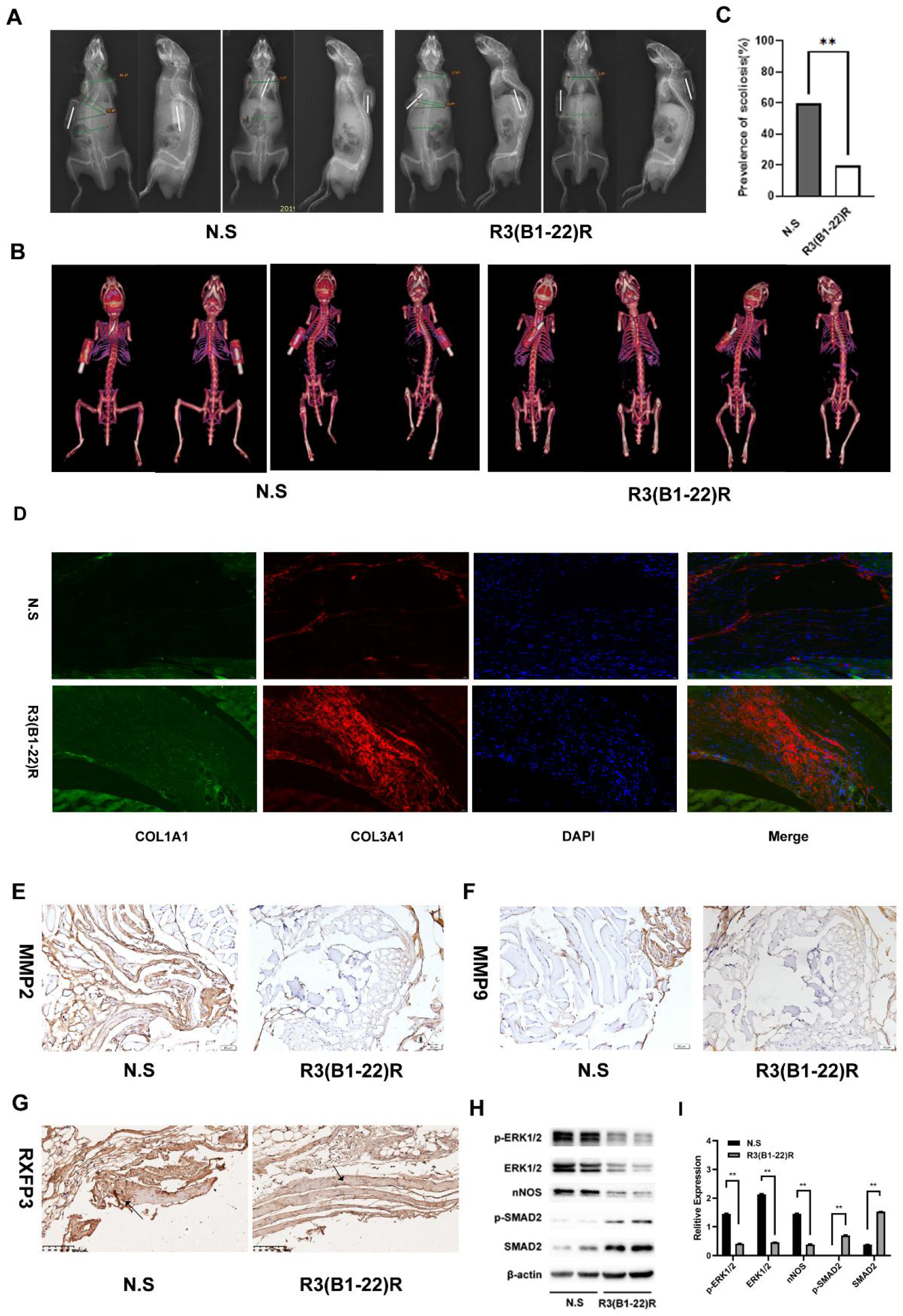
Inhibition of relxin-3 and RXFP3 by R3(B1-22)R prevented scoliosis in C57BL/6J mice. **A–B**, Scoliosis characterization in C57BL/6J mice was evaluated by X-ray and 3D-CT. **C**, The scoliosis incidence analysis between R3(B1-22)R-treated mice and NS-treated mice. The scoliosis incidence was significantly lower in R3(B1-22)R-treated mice than in NS-treated mice (22% vs. 62%). **p<0.01. **D**, The representative immunofluorescence image of COL1A1 (Green) and COL3A1 (Red) in ligament tissues from the R3(B1-22)R and NS groups. The increased expression of COL1A1 and COL3A1 was observed in spinal ligament tissues from the R3(B1-22)R group. DAPI (Blue) was used to stain the cell nuclei. **E–F**, The representative immunohistochemistry image of MMP2 and MMP9 in ligament tissues from the R3(B1-22)R and NS groups. Lower expression of MMP2 and MMP9 was detected in spinal ligaments from the R3(B1-22)R group, compared with those from the NS group. **G**, The representative immunohistochemistry image of RXFP3 in ligament tissues from the R3(B1-22)R and NS groups. No significant difference in RXFP3 expression was found in spinal ligament between R3(B1-22)R and NS groups. **H–I**, The expression of ERK1/2 and TGF-β/SMAD signaling in ligament tissues from the R3(B1-22)R and NS groups. Western blot assays confirmed lower expression of ERK1/2 and neuronal nitric oxide synthase in spinal ligaments from the R3(B1-22)R group (n=5), compared with those from the NS group (n=5). The expression of phosphorylated SMAD2 was significantly increased in spinal ligaments from the R3(B1-22)R group. The relative protein level was evaluated based on the gray value of the corresponding band. *p<0.05, and **p<0.01.

## Discussion and conclusion

Ligament laxity is observed in most AIS patients, and relaxin and relaxin-related peptides markedly contributes to the ligament laxity. However, correlation of relaxin and relaxin-related peptides and AIS is still poorly understood. In this study we found relaxin3 plasma level and RXFP3(relaxin family peptide receptor 3) expression was significantly higher in AIS patients and classic scoliosis mice model. Furthermore, R3(B1-22)R (RXFP3 antagonist) significantly decreased the prevalence of scoliosis in C57BL/6J mice scoliosis model. Our findings have uncovered that the anti-fibrotic effects of relaxin-3 and RXFP3 activation may aggravate the pathogenesis of AIS and provided a potential target treating to prevent this disease.

Until 1959, when scoliosis was induced in chickens via surgical removal of the pineal gland, there was no adequate AIS animal experimental models^(*26*)^. Melatonin, also known as N-acetyl-5-methoxytryptamine, is the major hormone secreted by the pineal gland. To demonstrate the relationship between melatonin and induction of scoliosis, an AA-NAT gene knock-out strain of mice with congenital melatonin deficiency (C57BL/6J) was developed^(*21*)^. Compared with 25% of quadrupedal mice, 70%–100% of melatonin-deficient bipedal C57BL/6J mice developed spinal deformity^(*27*)^. Another study indicates that C57BL/6J bipedal mice can serve as a steady scoliosis model in AIS research^(*28*)^ even though no significant change in melatonin levels has been observed in patients with AIS^(*29, 30*)^ and melatonin has not been shown to have therapeutic effects in scoliosis^(*31, 32*)^. However, melatonin can inhibit the hypothalamic-pituitary-gonadal axis, and its secretion reduction can promote the development of gonads^(*33, 34*)^. Thus, the secondary effect of melatonin, rather than the melatonin itself, may contribute to the etiology of AIS. Rln3 knockout C57BL/6J mouse model confirmed that RLN3 deficiency could lead to a significant decrease in scoliosis incidence rate in C57BL/6J scoliosis animal models, suggesting that RLN3 may play an important role in the pathogenesis of AIS. However, systemic knockout mice may have some disadvantages, such as complex phenotype and weak tissue specificity. We plan to use conditional knockout mice in later experiments to further improve our experimental study.

Relaxin peptide family is a recently found peptide family that is relate to ligament laxity^(*35*)^. In humans, the relaxin peptide family is encoded by seven genes: relaxin genes RLN1, RLN2(relaxin), and RLN3 and insulin-like peptide genes INSL3, INSL4, INSL5, and INSL6. Most other species have only five of these genes, including RLN1 and RLN3, which are the species equivalents of human RLN2 and RLN3, respectively^(*35*)^. The function of the RLN1 gene in humans and higher primates is unknown^(*36*)^. RLN2, the mammalian 6-kDa heterodimeric polypeptide hormone, is a member of the insulin-like superfamily and consists of seven peptides of high structural but low sequence similarity^(*37*)^. It was first described by Frederick Hisaw in 1926, who showed in animals that ligaments relaxed following injection with pregnant-guinea-pig plasma^(*36*)^. RLN2 is associated with gonadal development and correlated with estrogen levels through pulsed secretion from tissues, such as gonads^(*38, 39*)^. RLN2 also plays important roles in collagen catabolism of the pubic symphysis during gestation in lower mammals, such as mice and rats^(*40, 41*)^. RLN3 is the most recently identified relaxin family peptide^(*42*)^. It is included as a “relaxin” peptide because of its characteristic RxxxRxxI/V relaxin binding motif in the B-chain, but otherwise it has relatively low sequence homology to other relaxin peptides. Unlike other relaxins, the sequence of RLN3 is well conserved across species^(*43-46*)^. RLN3 also is believed to be the ancestral peptide of this family. In mammals, it is primarily a neuropeptide involved in stress, memory, and appetite regulation^(*42, 47*)^. So far, few has been understood on the role of RLN3 outside the nervous system. Our study showed that RLN3 plasma level may have potential to be early serological indicators for AIS screening. As an ancestral molecule, RLN3 may have a broader effect to be studied.

Relaxin can lead to ligament laxity by its anti-fibrotic effects. Elevated RLN2 serum levels can increase pelvic width and height in pregnant cattle^(*48*)^, as well as tendon^29^ and joint laxity^(*49-51*)^ in humans. The relaxin receptor RXFP1 has been found in human female anterior cruciate ligaments^(*52*)^. Ample research suggests that RLN2 can increase laxity of the anterior cruciate ligament^(*53, 54*)^, thus increasing risk of injury^(*54, 55*)^. RLN2 also changes the mechanics of the ligament via anti-fibrotic effects mediated by discharge of MMPs^(*56*)^, collagenase^(*57*)^, and plasminogen activator^(*58*)^. RLN3 may have an anti-fibrosis role as RLN2. RLN3 recently has been shown to promote MMP2 levels in a dose-dependent manner when administered to rat ventricular fibroblasts in vitro^(*59*)^. In this way, RLN3 behaves in a comparable manner to RLN2^(*18*)^. Moreover, RLN3 may enhance the collagen-inhibitory effects of RLN2^(*18*)^. This research confirmed that RLN3 induced an anti-fibrotic effect on both human and mice spinal ligament fibroblasts, indicating that RLN3 increased the incidence of scoliosis by elevating spinal ligament laxity through its collagenolytic effect. The ability of RLN to inhibit TGF-β1-mediated spinal fibrosis progression has been shown to involve activation of the MAPK-ERK1/2 pathways, downregulation of Smad2^(*60*)^, and phosphorylation. In this study, we found that ERK1/2 plays an important role in the process of RLN3 acting on ligament fibroblasts, although ERK1/2 is involved in many signaling pathways. This study is still a preliminary study. We are going to study each molecule of RLN3 through its receptor in the follow-up study to find out whether there are any key molecules to the pathogenesis of AIS.

RXFP3 antagonist may have potential for prophylactic treatment of AIS. The receptors for RLN3 include its endogenous receptor RXFP3, as well as RXFP1 and RXFP4 with high affinity. Previous studies have suggested that RXFP3 is generally expressed in brain system, while RXFP1 can be expressed in most of fibroblasts^(*25*)^. In our experiment, we found that RXFP3 can significantly expressed in spinal ligament fibroblasts, as well as RXFP1. As RXFP3 has the strongest affinity for RLN3^(*61*)^.RLN3 is likely to work by binding to RXFP3 in spinal ligament fibroblasts. Luckily there is plenty of antagonist of RXFP3 compared with RXFP1 which has few antagonist peptide (B-R13/17K, as well as partial agonist) by now. The R3(BD23–27)R/I5 is an RXFP3 antagonist that has been instrumental in defining the physiologic functions of the receptor. However, studies showed that it may only block some but not all pathway activated by RXFP3. R3(B1-22)R is another antagonist of RXFP3 with high affinity, and far easier to produce as single-chain peptide, furthermore it is specific for RXFP3. This study confirmed that R3(B1-22)R blocked the anti-fibrotic effect of RLN3 on spinal ligament fibroblasts and significantly reduced the incidence of scoliosis in a classic mouse model (C57BL/6J) of scoliosis. We observed overexpressed RXFP3 in ligaments fibroblast of AIS patients (Figure 1E, Figure 3D). Thus, RXFP3 antagonist such as R3(B1-22)R may have potential for prophylactic treatment of AIS.

In sum, AIS is a complex global spinal disease that occurs through multiple factors in multiple systems^(*62, 63*)^. Its pathogenesis remains unknown. The results of this study suggest that the anti-fibrotic effects of RLN3-RXFP3 activation may play an important role in the pathogenesis of AIS.

## Author contributions

Zhanghao, Luo beier, Sun fengyuan contributed equally to this work. Zhanghao and Sun fengyuan performed the experiment; Luo beier performed the data analyses and wrote the manuscript; Xu bing contributed significantly to analysis and manuscript preparation; Wang guokun and Ming Li helped perform the analysis with constructive discussions.

---

**Table 1.** General characters of subjects enrolled in the study

|  | AIS group | Control group | *P*-value |
| --- | --- | --- | --- |
| Age (years) | 12.0 ± 1.5 | 12.7 ± 1.8 | 0.18 |
| BMI (kg/m<sup>2</sup>) | 19.3 ± 1.5 | 18.8 ± 2.7 | 0.41 |
| Menarche (years) | 10.3 ± 0.9 | 10.6 ± 0.8 | 0.45 |
| Risser (°) | 2.0 ± 1.1 | 2.3 ± 1.0 | 0.20 |
| VAS(points) | 1.7 ± 0.9 | 0.6 ± 0.7 | 0.00 |
| CRP(mg/L) | 3.3 ± 1.0 | 3.2 ± 0.9 | 0.65 |
| ESR(mm/H) | 6.2 ± 1.9 | 5.9 ± 1.8 | 0.58 |
| Curve Magnitude (°) | 36.8 ± 10.9 | / | / |
| JHM (points) | 4.1 ± 1.6 | 2.2 ± 1.1 | 0.00 |

BMI: body mass index, VAS: Visual analogue scale, CRP: C-reactive protein ESR: erythrocyte sedimentation rate, JHM: Joint hypermobility

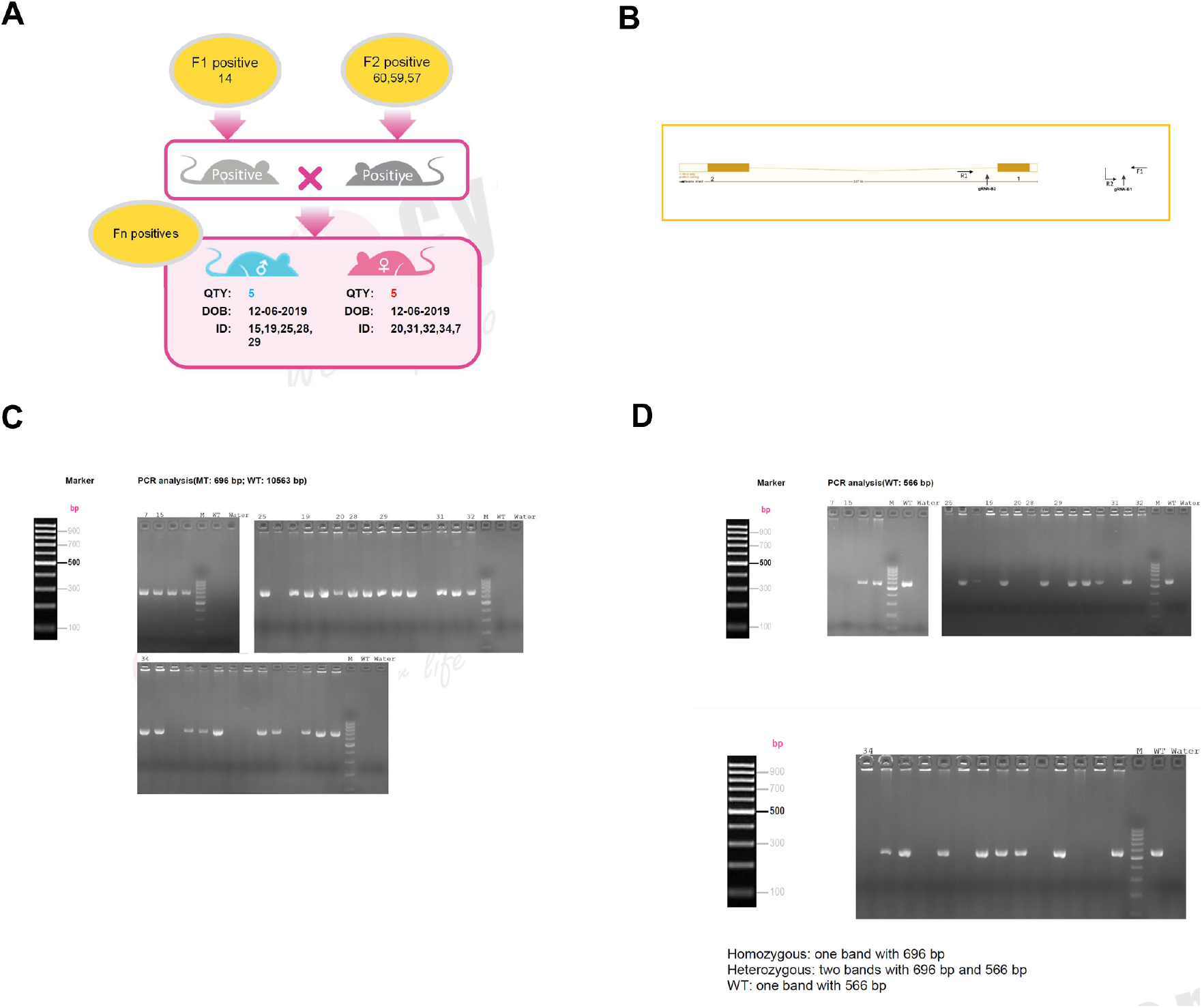

