## supplemental table1 for "The anti-fibrotic effects of RLN3 aggravated the pathogenesis of adolescent idiopathic scoliosis-a preliminary study"

**Supplement data**

**Fig. S1**

a C57BL/6J mouse model with RLN3 knockout (RLN3-/-) was constructed by the CRISPR/Cas9 system. A: Breeding Scheme: The gRNA to mouse Rln3 gene, and Cas9 mRNA were co-injected into fertilized mouse eggs to generate targeted knockout offspring. F0 founder animals were identified by PCR followed by sequence analysis, which were bred to wildtype mice to test germline transmission and F1 animal generation. Inter-cross heterozygous F1 mice and heterozygous F2 mice to generate homozygous Fn mice. B Genotyping Strategy.C PCR Screening: PCR Primers:(F1): 5'-GGAAAGCTAGGTGTGGACTCTGG-3';Reverse primer (R1): 5'-GTCTTAGGCTGTCCTTGAGTTCATG-3'; Results: Fn animals 15, 19, 25, 28, 29, 20, 31, 32, 34 and 7 were identified positive by PCR screening. D Results： Fn animals 15, 19, 25, 28, 29, 20, 31, 32, 34 and 7 were identified homozygotes by PCR screening.

**Table S1**

JHM prevalence was found significantly higher in AIS patients than control group (p=0.004).

Table S1 JHM prevalence was found significantly higher in AIS patients than control group

| | AIS group# | Control group# | Significance |
|---|---|---|---|
| RLN3 | 84.942±64.481 | 29.415±14.267 | 0.000* |
| RLN2 | 9.716±11.081 | 8.395±7.714 | 0.649 |
| RLN1 | 76.689±50.052 | 82.285±44.125 | 0.696 |
| JHM present | 12 (54.5%) | 3(9.1%) | 0.004* |

*. The mean difference was significant at the 0.05 level.

#. Plus-Minus values are means ±SD
